# Strategic citations for a fairer academic landscape

**DOI:** 10.1101/2025.08.06.668908

**Authors:** Miriam Beck, Pavanee Annasawmy, Deborah Birre, Michela Busana, Nicolas Casajus, Camille Coux, Clara Marino, Nicolas Mouquet, Lisa Nicvert, Brunno F. Oliveira, Cathleen Petit-Cailleux, Axelle Tortosa, Mithila Unkule, Chloe Vagnon, Devi Veytia

**Affiliations:** FRB-CESAB; FRB-CESAB, MARBEC; FRB-CESAB, INRAE-LESSEM; FRB-CESAB, Universite de Toulouse

## Abstract

Scientific publishing is increasingly dominated by for-profit journals, which attract prestige and submissions through high impact factors (IF). While some of these partly reinvest in research and dissemination and can be considered academia-friendly, non-profit journals - those that fully reinvest revenue into the academic community - often struggle for visibility despite promoting more equitable publishing models. Using citation data from over 70,000 publications in ecology and evolution, we show that citation practices are siloed: for-profit journals disproportionately cite other for-profit journals, academia-friendly journals preferentially cite other academia-friendly journals, and non-profit journals likewise favor citations to non-profit sources. This asymmetry structurally reinforces the IF advantage of for-profit journals simply because they are dominant in the publishing system. To address this inequity, we propose a soft-power, low-risk approach of “strategic citation”. By deliberately choosing to cite relevant articles from non-profit journals when multiple references would be equally valid, researchers can contribute to increasing those journals’ visibility and IF. This approach preserves scientific rigor and does not restrict publishing choices, but adds a layer of ethical intentionality to citation practices. Strategic citation offers a practical, actionable lever for researchers to promote a more balanced and ethical publishing system that complements broader structural reforms.

## Introduction

Academic journals play an essential role in the scientific process by ensuring that new findings are critically evaluated through peer review, widely disseminated, and serve as a foundation for future research. This role carries a significant social responsibility for journals and their governing bodies. Journals operate over a broad spectrum of publishing strategies, ranging from strictly for-profit to non-profit [1–5]. The landscape of academic publishing has undergone a significant transformation over recent decades, marked by a notable shift towards for-profit models [6–8]. In contrast, publishers that invest surplus funds to support scientific research or its dissemination have seen a relative decline [7]. For example, the five largest for-profit publishers accounted for 20% of publications in the 1970s, a figure that rose to 53% in 2013 [9]. This shift may have been catalysed by the growing reliance on journal-level metrics, particularly the impact factor (IF), as proxies for journal prestige and research quality. Originally designed to measure citation frequency, the IF is now often used to indicate journal excellence and visibility [10], potentially leading to increased submissions and citations.

The increasing prevalence of for-profit journals has two major implications for the academic system. First, a system dominated by for-profit publishing increases the financial stress on academic institutions, libraries and researchers, who are subjected to rising subscription costs. The widening gap between the actual costs of publication and the fees charged to access these publications, both through subscription models and article processing charges, often far exceeds what could reasonably be considered fair or proportional compensation for reviewing and publishing services [11–13]. Second, the business models adopted by many for-profit journals have drawn considerable criticism from the research community and have been accused of prioritising financial gain over the equitable dissemination of knowledge [14–16]. This dynamic poses an existential challenge to the integrity of scientific research [17,18].

Academic researchers are pressured to publish in journals perceived as most excellent, a perception still largely based on IF, to the benefit of their career and institutions [19]. This pressure is particularly acute for early career researchers, who often face employment insecurity and rely heavily on their publication records to secure funding, fellowships, and academic appointments [20]. As a result, science, long upheld as a public good intended to serve the advancement of society [21,22], is arguably becoming a “tragedy of the commons” [23], where the pursuit of individual or institutional gain undermines the collective benefits of open, equitable knowledge production [24]. For-profit publishers, such as Springer, Reed-Elsevier, Wiley-Blackwell, and Taylor & Francis, now control an increasing share of high-IF journals [25]. Given the commercial logic underpinning for-profit journals and the central role of the IF in shaping perceptions of prestige and visibility, these journals are naturally driven to maximize their IF. This dynamic feeds a self-reinforcing cycle in which for-profit journals gain increasing visibility and submissions, further entrenching the dominance of the for-profit model (Figure 1). Notable exceptions challenge the narrative that only for-profit journals can succeed and demonstrate that non-profit journals can rise to the top of their fields. For instance, *Science* – a non-profit journal published by the American Association for the Advancement of Science – consistently ranks among the highest IF journals in the science and technology fields (IF ≈ 45 in 2024, [26,27]).

**Figure 1.**
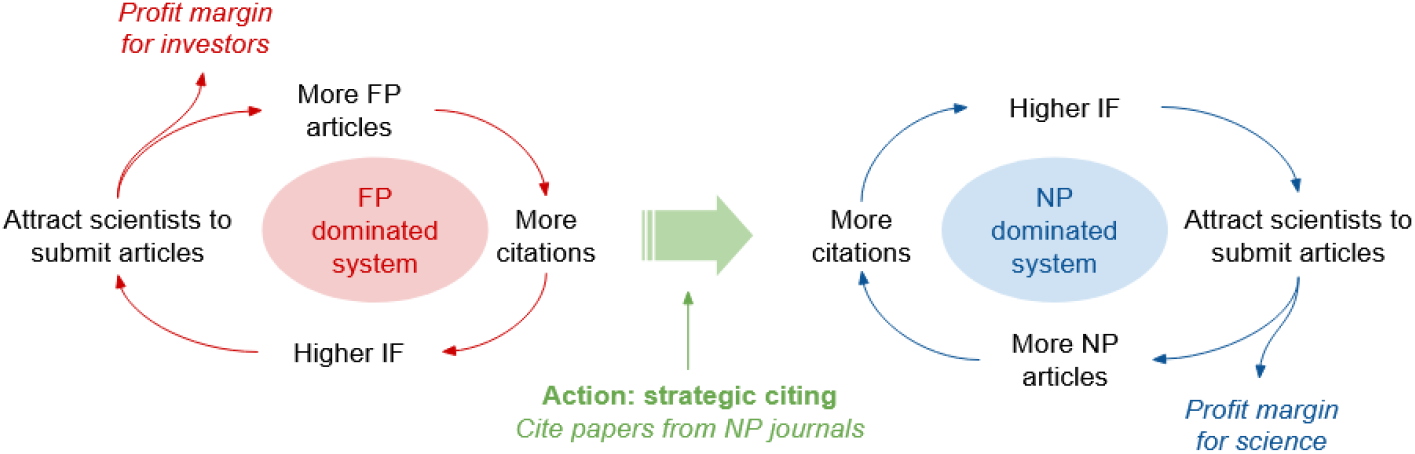
Illustration of a transition from a vicious to a virtuous circle of journal prestige. Strategic citing of articles from non-profit journals can help increase their impact factor, thus enhancing the perceived prestige and notoriety of non-profit journals. IF: impact factor; FP: for-profit journals; NP: non-profit journals.

However, the publishing landscape is more nuanced, and some journals published by for-profit companies reinvest revenues into research or dissemination, and can be considered “academia-friendly” [1]. These include journals affiliated with public research institutions or scholarly societies. For example, *Landscape Ecology* is published by Springer but is affiliated with the International Association for Landscape Ecology [26]. A more balanced system, where non-profit publishers also thrive, would benefit the academic and civil societies. Here we propose a constructive approach to shift the publishing landscape from a “vicious” to a “virtuous” cycle (Figure 1) by re-evaluating the citation practices of publishing researchers. By strategically choosing to cite articles published in non-profit journals, researchers can help enhance the visibility of these journals, and contribute to increasing the journals’ perceived prestige. Importantly, scientific quality and relevance must remain the primary criteria for citation, and any shift in citation behavior should happen within the bounds of scholarly norms grounded in content relevance and scientific rigor. In order to implement this strategy, we encourage authors to select sources published in non-profit or academia-friendly journals where multiple suitable references exist, such as for broadly accepted statements often found in the introduction or discussion sections of scientific publications (*i*.*e*., “The biodiversity crisis is accelerating worldwide”).

As a necessary first step towards this goal, we assessed the current state of citation patterns by quantifying the representation of distinct business models in the references of the scientific literature, exemplified for the fields of ecology and evolution. For these fields, an assessment of the journal’s business model has been facilitated *via* the DAFNEE database [1] (Database of Academia Friendly jourNals in Ecology and Evolution, https://dafnee.isem-evolution.fr/). The 2025 version of DAFNEE catalogues 611 journals that are oriented towards the academic community (i.e., “academia-friendly”) based on indicators like ownership, fee structure, institutional partnerships, and reinvestment practices. Based on this database, we analysed 70,848 articles published in 2023 across 270 journals following three different business models (Box 1). For each publication, we extracted the cited references and classified them based on the publishing journal as (a) for-profit, (b) for-profit academia-friendly, or (c) non-profit citations.

### Box 1.

**Characterizing current citation practices in ecology and evolution across journal business models**

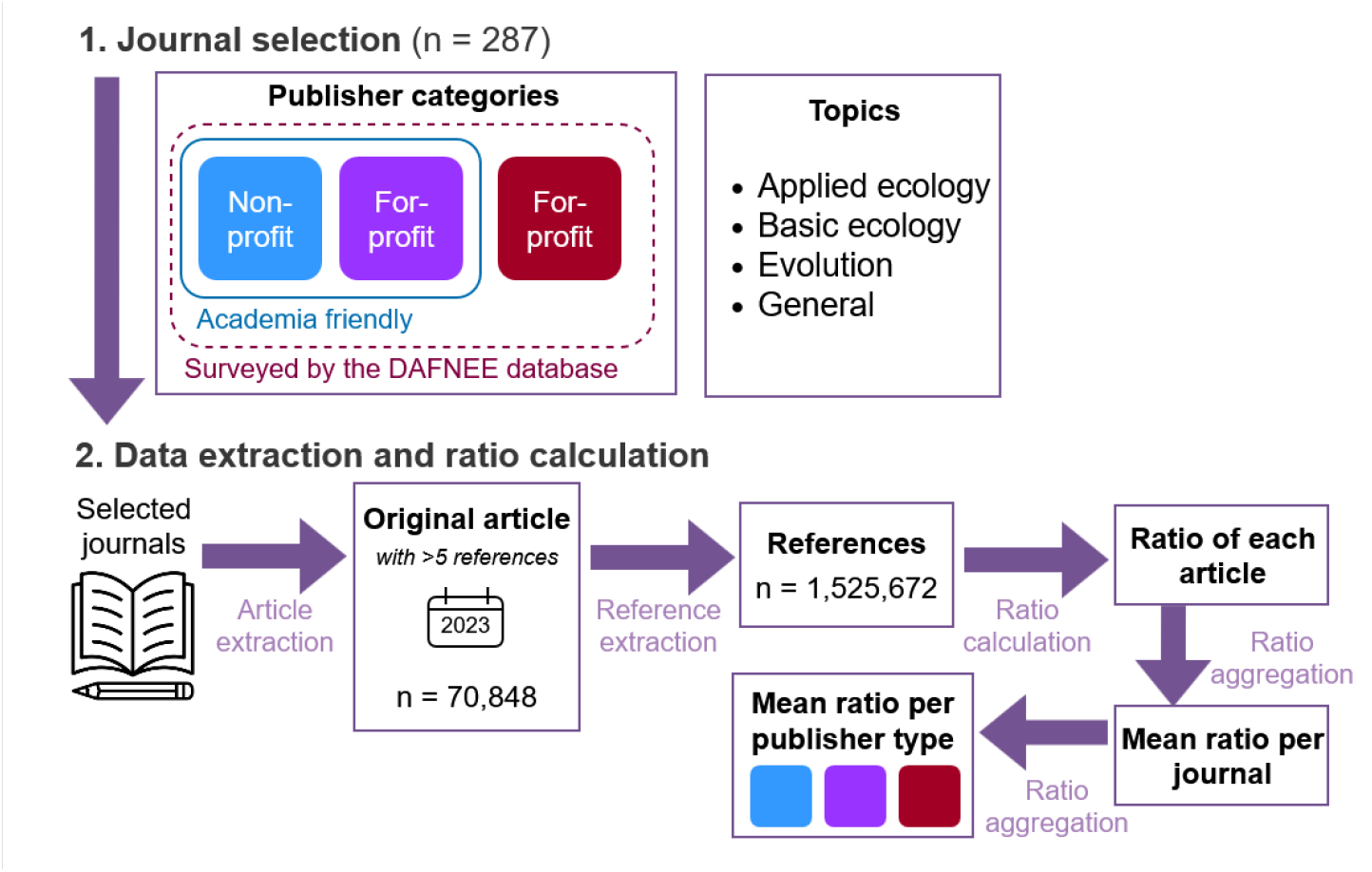

Our analysis is based on the DAFNEE database [1] that lists journals classified as “academia-friendly” (see figure above). We limited our analysis to journals within the thematic fields of “applied ecology”, “basic ecology”, “evolution” and “general” to minimize the variation in citation practices across disciplines. To represent the broader publishing landscape we include purely for-profit journals from the list of surveyed journals - evaluated by DAFNEE but not meeting the criteria for “academia friendly” - that matched our selected thematic fields. The business model of those journals was obtained from each journal’s website, while for academia-friendly journals it was retrieved from the DAFNEE database. The final journal list comprised 287 journals, among which 93 were classified as “non-profit”, 83 “for-profit academia-friendly” and 111 as “for-profit” (see S7).

Using OpenAlexR [28], we identified all articles published in 2023 by the selected journals (n = 130,324 “original articles”) and retrieved their cited references (n = 6,604,473). Each reference was classified based on the business model of its publishing journal into one of the three categories. Only references to journals in our selection were included in the further analysis. To reduce noise from atypical citation behavior and small sample sizes, we excluded publications with fewer than five references and removed special article types (*e*.*g*., comments, responses). This resulted in a final dataset of 70,848 original articles with 1,525,672 cited references (mean = 21.53 +/-18.01 SD references per article) from 270 journals. This filtering did not influence the general outcome (S6).

For each article, we calculated the proportion of its references published in each of the three categories of journals.These ratios were aggregated at the journal-level, and subsequently at the business model level, by computing unweighted mean citation ratios for each group. A more detailed description of the methods is provided in the supplementary material.

## Citation practices are siloed between for-profit and non-profit journals

If citation practices were completely unbiased, we would expect no significant differences in the proportion of citations towards for-profit *vs*. for-profit academia-friendly *vs*. non-profit journals. Instead, we found that publications in purely for-profit journals included significantly more references from for-profit journals than articles published elsewhere (Fig. 2; Dunn test with p < 0.001). Likewise, publications in for-profit but academia-friendly journals most frequently cited for-profit academia-friendly references (Fig. 2; Dunn test with p < 0.001), while publications in non-profit journals showed the highest proportion towards non-profit references (Fig. 2; Dunn test with p < 0.001). This suggests a strong tendency for researchers to cite articles published in journals within the same business model. In part, this may reflect a tendency to cite articles from the same journal (see S5), potentially to increase the chances of publication acceptance [29,30]. Beyond this, financial aspects might also play a role, since academia-friendly journals often have lower processing fees [1] . Researchers’ citation behaviour may also reflect their ethical views on the role of profit in academic publishing. For example, authors who choose to publish in non-profit journals may be more motivated to support the non-profit publishing system through their citation choices compared to authors publishing in for-profit journals. Another mechanism might be field-specific clustering, where certain subfields are predominantly covered by specific business models, thereby reinforcing internal citation loops within each business model.

**Figure 2.**
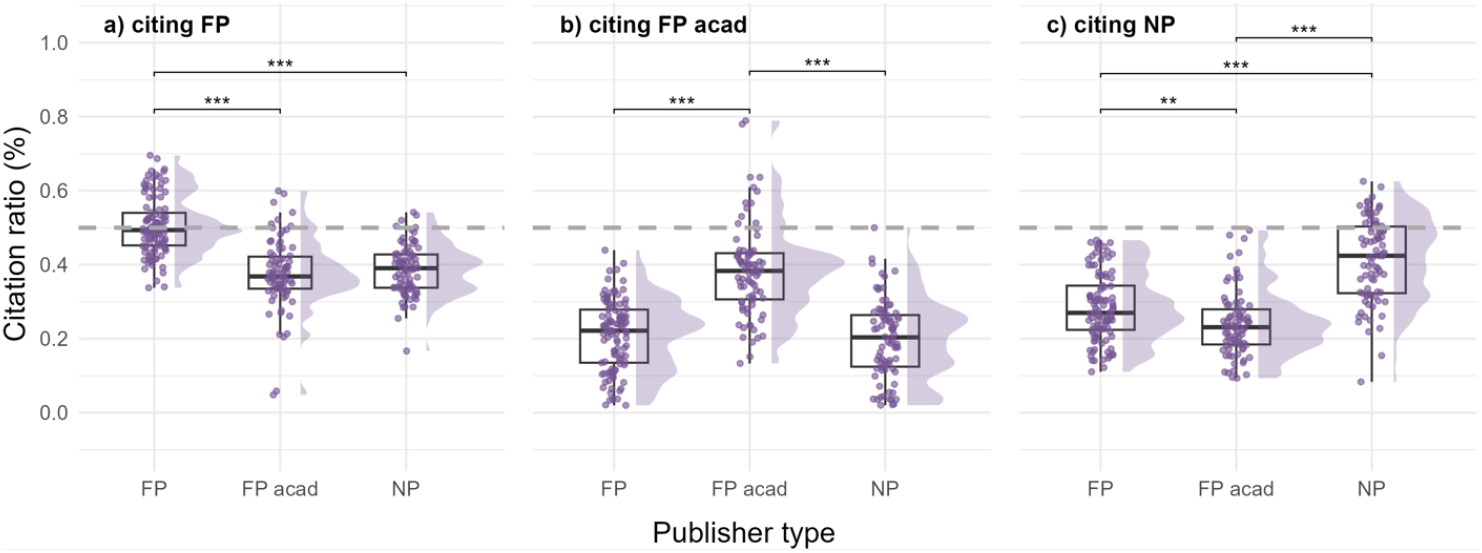
Proportion of citations towards articles published in (a) for-profit (FP), (b) for-profit academia-friendly (FP acad) and (c) non-profit (NP) journals among the same three publisher types. The boxplot indicates the median with the box comprising values between first and third quartile; scatters and density curve indicate the distribution of the datapoints. Horizontal brackets indicate statistically significant differences in citation ratio between two publisher types. Asterisks denote significance levels: p < 0.05 (*), p < 0.01 (**) and p < 0.001 (***), based on Dunn tests with Bonferroni correction following significant Kruskal-Wallis tests (p < 0.001). Non-significance is not indicated. The analysis is based on n=106, n=82 and n=82 journals for journals following for-profit, for-profit academia-friendly and non-profit business models, respectively.

Our results come with inherent limitations, largely due to possible bias from our focus on the journals listed or surveyed in the DAFNEE database. Every scientific field has its own publishing system, which may differ substantially from those observed in ecology and evolution. Moreover, the distinction between for-profit, and non-profit journals, even when considering in-between business types, may not always be obvious. A more thorough evaluation of citation practices would require including journals from a broader range of scientific fields to assess the robustness of our findings and to explore how these patterns vary across academic fields. In addition, a time series analysis could reveal whether citation practices have shifted over time, which would be particularly relevant given the rapid expansion of for-profit publishing in recent years. For instance, the portfolio of Nature-branded journals (published by Springer Nature, a for-profit publisher) has grown from a single flagship journal in 2009 to 34 titles in 2024 (see [7]), illustrating the accelerating commercialisation of scientific publishing.

## Strategic citation to support a balanced publishing system

Our results show a clear bias in citation practices: researchers tend to cite more articles published in journals that share the same business model (Fig. 1). Hypothetically, if both for-profit and non-profit journals received a similar boost from these within-group citations, their impact factors could grow at comparable rates. But in reality, the system is heavily imbalanced, with nearly five times more for-profit journals than non-profit ones [9]. As a consequence, purely by numbers, for-profit journals are more likely to accumulate citations, leading to faster growth in their IFs.

Even if non-profit journals maintain strong internal citation networks, the dominance of for-profit journals in the publishing landscape gives them a structural advantage in visibility and prestige. Although our analysis focuses on journals in ecology and evolution, similar patterns likely exist in other fields. Without efforts to correct this imbalance, there is a real risk that non-profit journals will continue to see their IF, and thus their visibility and attractiveness, decline over time. Few flagship titles, such as *Science*, might be able to retain their standing, while most will be overshadowed by a growing landscape of for-profit journals with stronger citation performance and perceived prestige.

For individual researchers who wish to promote non-profit journals, the most straight-forward action is to prioritise publishing in non-profit journals, or even to boycott for-profit journals [31]. However, such a strategy comes at significant cost, as choosing to publish exclusively in non-profit journals can significantly limit available publishing venues and access to high impact journals. Given that IF remains a widely used - albeit contested [10] - proxy for academic excellence, boycotting for-profit journals creates disproportionate risks for early-career researchers [20,32,33]. An alternative is to propose actions that meaningfully challenge the *status quo* while minimising personal risks [20]. While we acknowledge the validity of boycott approaches, we propose a soft leverage: a strategic choice of citations that promotes greater equity in academic publishing without requiring individual restrictions on the choice of venues. Although many for-profit journals are important active actors of the publishing landscape, we need to foster a healthier, more balanced system – one characterized by a fairer distribution of prestige and influence and not guided by profit margins.

What does this mean for action? The strategic citation approach we propose offers a practical way for researchers to support a more equitable publishing landscape. Scientific articles are typically structured around a review of the state of the art in the introduction and often include general statements supported by well-known references. For example, in biodiversity science, authors might cite studies on the global extinction crisis, while in climate research, references to global warming trends are common. Similar types of general or widely accepted claims also appear in the discussion or perspective sections. In these cases, where multiple valid sources exist, authors can choose to cite relevant articles published in non-profit journals, if available. This approach does not compromise scientific accuracy but adds a layer of intentionality to citation choices. These choices could be implemented as an additional step during manuscript preparation: after the content is finalised, authors could calculate the ratio of for-profit *vs*. non-profit references and, where possible, adjust it. Accompanying this article, we provide the easy-to-use R package “fairpub” ([removed for peer-review]) that allows users to quantify the proportion of non-profit and/or academia-friendly references based on a BibTeX file (currently limited to the fields of ecology and evolution). Given our finding that articles published in for-profit journals tend to strongly favour citations from other for-profit journals (and *vice versa* for non-profit and academia-friendly publications), this check should be prioritised by authors publishing in for-profit journals. We concur with previous studies that emphasise scientific quality and relevance must remain the primary criteria for citation choice (*i*.*e*., [34]). However, with most articles containing dozens of references, there is meaningful room to shift citation patterns without compromising scientific rigor. Ultimately, this approach will not replace for-profit journals, but maintain the visibility and perceived quality of non-profit journals while the academic community works toward broader reforms. Our proposal is a complementary, low-risk strategy that can help sustain a balanced publishing system during this necessary transition.

To be truly effective, researchers need awareness of the issue and access to reliable, transparent information about journal business models, something that is not currently easily accessible. While initiatives like the DAFNEE database and the “fairpub” package can provide valuable insights for ecology and evolution journals, similar tools are largely missing in other disciplines. Moreover, as business models evolve and ownership structures shift, these databases must be regularly updated. Ideally, a neutral regulatory body or institution could evaluate and clearly label journals based on how their revenues are used. Greater transparency and awareness will enable researchers to make informed, strategic publishing and citations choices and foster positive change across the publishing system toward a fairer academic landscape

## Conclusion

The current academic publishing system is structurally imbalanced, with for-profit publishers dominating publication volume and increasing their perceived excellence [35] through IF inflation. Their reliance on subscriptions and article processing charges shifts financial burdens onto researchers and institutions, reinforcing global inequalities in access and visibility [36,37]. Non-profit journals offer more equitable models, such as diamond open access (*i*.*e*., no publication nor reading fees), but remain under-resourced in a system driven by the for-profit vicious circle (Figure 1) [38]. While a complete transition to non-profit publishing is unrealistic under the market-oriented systems prevalent in many leading research nations, reinforcing the visibility and legitimacy of non-profit journals can incrementally boost their IF, helping them (re)gain recognition and compete within the current evaluation systems. There is still time to change and sustain the virtuous cycle of non-profit and for-profit, academia-friendly journals (Figure 1), provided we have reliable information on journals’ business models. In the short-term, the commitment to research ethics should extend to how we cite, because it begins with the choices we make in the reference lists. This individual-level action complements long-term, broader institutional reforms needed to reduce reliance on journal-level metrics and re-establishing science as a public good.

## Code and data availability

All data and code needed to evaluate the conclusions in the paper are present in the corresponding git repository: [removed for peer-review]. This paper cites 60% NP journals and 40% FP journals.

