## Supplementary material for "Strategic citations for a fairer academic landscape"

#### Materials and methods - Details

To characterize the landscape of citations in ecology and evolution, we retrieved all papers published in a selection of academic journals (see S7) for the year 2023. From each of these published articles, we retrieved all cited references to compute its citation ratio (*i.e.* the proportion of references of articles published in for-profit, for-profit academia-friendly and non-profit journals). Finally, we compared the citation ratio between journals with distinct publishing models.

##### Tools and selection of journal publishing model data

We used the DAFNEE (Database of Academia Friendly jourNals in Ecology and Evolution, https://dafnee.isem-evolution.fr/) database to obtain a list of academic journals and their associated business models. The DAFNEE initiative represents an initial effort to centralise information regarding the business models of journals in the fields of ecology and evolution. As of July 1st 2025, a total of 1,169 journals have been surveyed. To be included in the database, on top of criteria such as peer-review and international scope, journals need to be 'academia-friendly', *i.e.* a criteria based on the existence of a visible partnership with an academic institution or a non-profit organization. For the >611 journals that were classified as academia-friendly, DAFNEE also informs on whether a journal publisher operates on a non-profit basis (NP), allocates part of its profits to science by supporting or partnering with an academic institution (university press), or is fully for-profit (FP). For our analysis, the university press category was merged into the NP category because we considered that the benefit would be used to serve the scientific community through the university. From these 611 academia-friendly journals, we kept 172 journals in the fields of 'applied ecology', 'basic ecology', 'evolution' and 'general'. Among the remaining international scope journals that were surveyed but deemed non-academia-friendly (and therefore not listed in the DAFNEE database), we selected those relevant to the same categories to represent the whole variety of journal landscape (n=128). The publisher type of these journals was not provided by DAFNEE and was manually obtained based on information on the journal's website. The few journals identified as 'non-profit' generally belonged to publishers linked to the scientific community (n=13) and were thus integrated into the category of non-profit journals.

To retrieve articles and their cited references, we used the OpenAlex bibliographic database (Priem et al. 2022). OpenAlex is an open-access database of scholarly publications and a reliable alternative to the proprietary bibliometric databases (such as Web of Science and Scopus, Culbert *et al.* 2025). Furthermore, it is well suited for reproducible research: we used the R package openalexR (Aria *et al.* 2024) to query their API and retrieve article metadata.

##### Article and reference extraction

Among the 300 initial academic journals we sampled, only 287 were indexed in the OpenAlex database. We limited our analysis to 2023 and extracted metadata of all articles published in these 287 academic journals during that year. Only 270 academic journals published articles in 2023 with a total of 130,324 articles (median = 103 articles per journal, mean = 475.64 +/- 1848.25 SD). Finally, from each article, we extracted all cited references (n = 6,604,473, median = 48, mean = 50.68 +/- 33.77 SD per article).

#### Citation ratio computation

We matched each cited reference with its journal's publishing model (FP, FP-academia-friendly, NP). As we did only have this information available for our selected journals list, the number of articles with covered references (*i.e.*, references citing works that were among our list of journals) decreased to 110,988 and the number of cited references decreased to 1,633,350. In order to reduce biases due to a low number of cited references and to remove special article types (*i.e.*, comment, response), we excluded articles with less than 5 cited references. After applying this filter, our data comprised 70,848 articles with 1,525,672 cited references (mean = 21.53 +/- 18.01 SD references per article) from 270 journals (S1). Including articles citing <5 references did not affect the overall finding (S6).

| | For-profit | For-profit academia friendly | Non-profit | |
|---|---|---|---|---|
| **Journals** | 106 | 82 | 82 | [illegible] |
| **Articles** | 43,285 | 10,661 | 16,902 | [illegible] |
| **Cited references** | 901,020 | 281,281 | 343,371 | [illegible] |

***S1.*** *Number of journals, articles, and cited references after filtering used to compute citation ratio per publisher type.*

For each article, we calculated the ratio of the three categories of cited references, considering only those for which publisher type information was available (as described above). Then we calculated the journal-level ratio by averaging ratios of all articles per journal. The distributions of these citation ratios are presented in Fig. 2 in the main manuscript. Finally we averaged these journal-level ratios within publisher type (S2).

Differences in citation practices between journals following distinct publisher types were tested for significance by Wilcoxon tests, followed by Dunn Test. P-values were adjusted for multiple comparisons following the 'Bonferroni' method (Dunn, 1961).

| | | Citation ratio | | |
|---|---|---|---|---|
| | | **For-profit** | **For-profit academia-friendly** | **Non-profit** |
| **Publisher type** | **For-profit** | 0.50 +/- 0.08 | 0.21 +/- 0.09 | 0.28 +/- 0.09 |
| | **For-profit academia friendly** | 0.38 +/- 0.09 | 0.38 +/- 0.12 | 0.24 +/- 0.09 |
| | **Non-profit** | 0.39 +/- 0.07 | 0.20 +/- 0.11 | 0.41 +/- 0.11 |

***S2.*** *Mean proportion and standard deviation of for-profit, for-profit (academia-friendly) and non-profit references in journals published by the three types of publishers.*

#### Sensitivity of results

##### Sensitivity towards self-citation

We repeated the calculation and aggregation of citation ratios after removing self-citing references, *i.e.* references published in the same journal as the original article. This reduced the number of articles and cited references (S3) to be considered and the effect size (S4, S5).

| | For-profit | For-profit academia friendly | Non-profit | Total |
|---|---|---|---|---|
| **Journals** | 106 | 82 | 82 | 270 |
| **Articles** | 38,015 | 8,620 | 14,786 | 61,421 |
| **Cited references** | 777,685 | 236,572 | 289,979 | 1,304,236 |

***S3.*** *Number of journals, articles, and cited references after filtering used to compute citation ratio per publisher type after removing self citation*

| | | Citation ratio | | |
|---|---|---|---|---|
| | | **For-profit** | **For-profit academia-friendly** | **Non-profit** |
| **Publisher type** | **For-profit** | 0.45 +/- 0.07 | 0.24 +/- 0.10 | 0.32 +/- 0.11 |
| | **For-profit academia friendly** | 0.43 +/- 0.09 | 0.28 +/- 0.12 | 0.28 +/- 0.12 |
| | **Non-profit** | 0.43 +/- 0.07 | 0.22 +/- 0.12 | 0.34 +/- 0.11 |

***S4.*** *Mean proportion and standard deviation of for-profit, for-profit (academia-friendly) and non-profit references in journals published by the three types of publishers after removing self citation.*

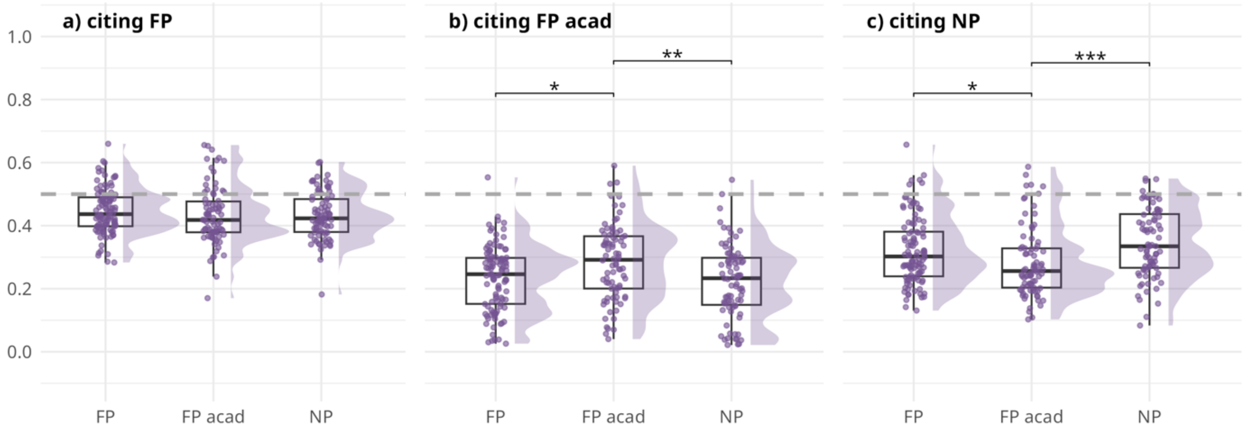


***S5:*** *Proportion of citations towards articles published in for-profit (a) for-profit (academic-friendly) (b) and non-profit journals (c ) among the same three publisher types after removing self-citation, i.e. articles citing the journal they are publishing in.*

Sensitivity towards filtering steps

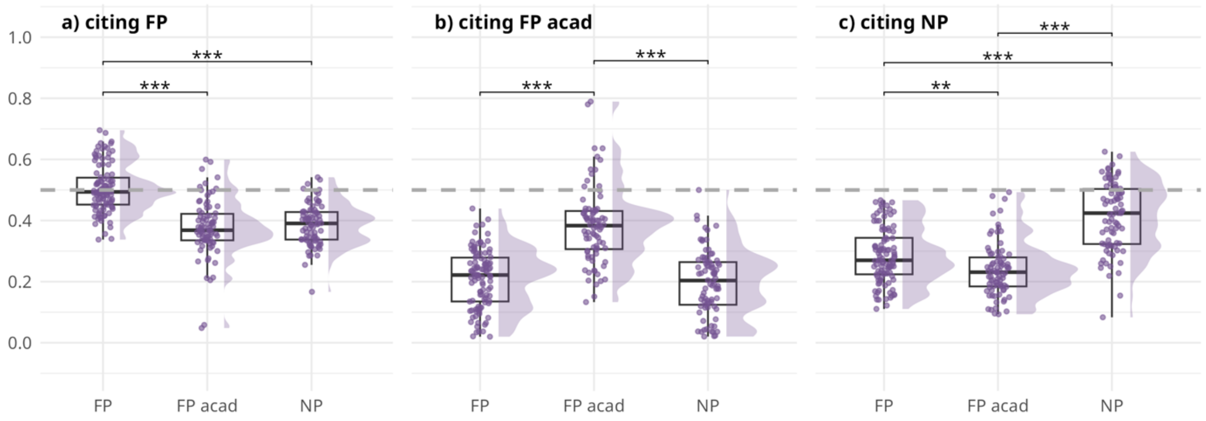


***S6:*** *Proportion of citations towards articles published in for-profit (a) for-profit (academic-friendly) (b) and non-profit journals (c ) among the same three publisher types including articles with <5 references.*

### Code availability

The code to reproduce data extraction and analysis is available on GitHub: [removed for peer-review]

***S7****. List of 270 journals included in the analysis that published articles in 2023 and their classification into business type categories*

| For-Profit | For-Profit (academia-friendly) | Non-Profit |
|---|---|---|
| acta ethologica | African Journal of Ecology | Acta Amazonica |
| Acta Oecologica | African Journal of Range and Forage Science | African Journal of Wildlife Research |
| Applied Soil Ecology | Agronomy for Sustainable Development | Animal Biodiversity and Conservation |
| Aquaculture Environment Interactions | AMBIO | Annual Review of Ecology Evolution and Systematics |
| Aquatic Botany | Animal Behaviour | Antarctic Science |
| Aquatic Conservation Marine and Freshwater Ecosystems | Animal Conservation | Aquatic Ecosystem Health & Management |
| Aquatic Ecology | Annals of Forest Science | Behavioral Ecology |
| Behavioral Ecology and Sociobiology | Annals of the New York Academy of Sciences | Biological Journal of the Linnean Society |
| Behaviour | Applied Vegetation Science | Biology Letters |
| Biodiversity and Conservation | Aquatic Living Resources | Biology Open |
| Biodiversity Data Journal | Archives of Sexual Behavior | BioScience |
| Biological Conservation | Austral Ecology | Bird Conservation International |
| Biological Invasions | Basic and Applied Ecology | Bulletin of Entomological Research |
| BMC Ecology and Evolution | Behavior Genetics | Bulletin of Marine Science |
| BMC Zoology | Biological reviews/Biological reviews of the Cambridge Philosophical Society | Canadian Journal of Fisheries and Aquatic Sciences |
| Communications Biology | Biotropica | Canadian Journal of Forest Research |
| Crustaceana | Bulletin of the Ecological Society of America | Cities and the Environment |
| Current Biology | Chelonian Conservation and Biology | Cold Spring Harbor Perspectives in Biology |
| Current Opinion in Insect Science | Cladistics | Comptes Rendus Biologies |
| Diversity | Coastal Management | Conservation Physiology |
| Diversity and Distributions | Conservation and Society | Ecology and Society |
| Ecological Complexity | Conservation Biology | eLife |
| Ecological Indicators | Conservation Letters | Environmental Conservation |
| Ecological Informatics | Conservation Science and Practice | European Journal of Taxonomy |
| Ecological Modelling | Ecography | Evolution |
| Ecology and Evolution | Ecological Applications | Evolution Letters |
| Ecology Of Freshwater Fish | Ecological Monographs | Evolutionary Journal of the Linnean Society |
| Ecosystems | Ecological Research | Freshwater Science |
| Endangered Species Research | Ecological Solutions and Evidence | Frontiers of Biogeography |
| Environmental Biology of Fishes | Ecology | Gayana |
| Environmental DNA | Ecology Letters | ICES Journal of Marine Science |
| Environmental Microbiology | Ecoscience | Ideas in Ecology and Evolution |
| Environmental Microbiology Reports | Ecosphere | Integrative and Comparative Biology |

European Journal of Wildlife Research
EvoDevo
Evolution & Development
Evolutionary Applications
Evolutionary Biology
Evolutionary Ecology
Fire
Food Webs
Forest Ecology and Management
Forests
Freshwater Biology
Frontiers in Ecology and Evolution
Frontiers in Marine Science
Frontiers in Microbiology
Global Change Biology
Global Ecology and Biogeography
Global Ecology and Conservation
Hydrobiologia
IAWA Journal
Insects
International Journal of Fisheries and Aquatic Studies
iScience
Journal for Nature Conservation
Journal of Applied Entomology
Journal of Arid Environments
Journal of Biogeography
Journal of Environmental Management
Journal of Experimental Marine Biology and Ecology
Journal of Forestry Research
Environmental Science and Pollution Research
Environmental Toxicology and Chemistry
Estuarine Coastal and Shelf Science
European Journal of Soil Science
Evolutionary Systematics
Frontiers in Ecology and the Environment
Functional Ecology
Hormones and Behavior
International Journal of Remote Sensing
Journal of Applied Ecology
Journal of Arid Land
Journal of Chemical Ecology
Journal of Coastal Conservation
Journal of Ecohydraulics
Journal of Ecology
Journal of Evolutionary Biology
Journal of Limnology
Journal of Mammalian Evolution
Journal of Sustainable Agriculture and Environment
Journal of Systematics and Evolution
Journal of the Marine Biological Association of the United Kingdom
Journal of Vegetation Science
Journal of Wildlife Management
Knowledge and Management of Aquatic Ecosystems
Landscape Ecology
Management of Biological Invasions
Marine Ecology Progress Series
NeoBiota
New Phytologist
Interface Focus
International Journal of Plant Sciences
International Journal of Wildland Fire
Invasive Plant Science and Management
Invertebrate Systematics
Journal of Coastal Research
Journal of Economic Entomology
Journal of Experimental Biology
Journal of Pollination Ecology
Journal of the National Museum (Prague) Natural History Series
Journal of Threatened Taxa
Journal of Tropical Ecology
Marine and Freshwater Research
Mediterranean Marine Science
National Science Review
Natural Areas Journal
New Zealand Journal of Ecology
Northwestern Naturalist
Novitates Caribaea
Open Biology
Oryx
Pacific Conservation Biology
Parasitology
Peer Community In Ecology
Peer Community in Evolutionary Biology
Peer Community In Forest and Wood Sciences
Peer Community In Registered Reports
Philosophical Transactions of the Royal Society B Biological Sciences
Philosophy Theory and Practice in Biology

Journal of Land Use Science
Journal of Pest Science
Journal of Sea Research
Journal of Theoretical Biology
Journal of Zoological Systematics & Evolutionary Research
Land
Limnologica
Mammal Research
Mammalia
Marine Biodiversity
Marine Biology
Marine Biology Research
Microbial Ecology
Molecular Ecology
Molecular Ecology Resources
Molecular Phylogenetics and Evolution
Movement Ecology
Mycorrhiza
Nature
Nature Communications
Nature Ecology & Evolution
Nature Plants
Nature Sustainability
Neotropical Biodiversity
Nova Hedwigia
Oecologia
One Earth
Open Journal of Ecology
Parasitology Research
PeerJ
Perspectives in Plant Ecology Evolution and Systematics
Phytotaxa
Plant Systematics and Evolution
Review of Palaeobotany and Palynology
Reviews in Fish Biology and Fisheries
Scientific Data
Scientific Reports
Studies on Neotropical Fauna and Environment
Thalassas An International Journal of Marine Sciences
The Botanical Review
Oikos
Open Research Europe
Organisms Diversity & Evolution
Pedosphere
People and Nature
Personality and Individual Differences
Perspectives in Ecology and Conservation
Population Ecology
Remote Sensing in Ecology and Conservation
Restoration Ecology
Taxon
Tropical Ecology
Urban Ecosystems
Vegetation Classification and Survey
Weed Research
Wellcome Open Research
Wetlands
Wildlife Biology
Wildlife Society Bulletin
Zoosystematics and Evolution
Phytobiomes Journal
Plant Ecology and Evolution
PLoS Biology
PLoS ONE
PNAS Nexus
Polish Journal of Ecology
Proceedings of the National Academy of Sciences
Proceedings of the Royal Society B Biological Sciences
Records of the Australian Museum
Royal Society Open Science
Science
Science Advances
Scientia Marina
Silva Balcanica
Sociobiology
Systematic Biology
The American Naturalist
The Bulletin of zoological nomenclature
The Coleopterists Bulletin
Web Ecology
Wildlife Research
Zoological Journal of the Linnean Society
Zoosystema

The Science of Nature
Theoretical Ecology
Theoretical Population Biology
Trends in Ecology & Evolution
Tropical Conservation Science
